# Analysis of the current state of frailty indexes and their implementation for aging intervention studies

**DOI:** 10.1101/2025.02.24.639506

**Authors:** Oliver G Frost, Anna Barkovskaya, Michael Rae, Abdelhadi Rebbaa, Amit Sharma

## Abstract

Animal lifespan studies are foundational to developing interventions against the biological aging process. In recent years, there has been rising interest in characterizing the effects of longevity therapeutics on health span. Frailty indexes, originally developed to assess clinical frailty in aging humans, have shown promise as measurements of biological age and have been adopted for use in rodent aging biology. This perspective looks at the current state of rodent frailty indexes and how they are implemented. The differences in frailty parameters used to calculate these indexes have led to inconsistencies between studies defining frailty. In this perspective, we have highlighted those differences and made recommendations for implementing protocols for frailty index measurement.

## 1. Introduction

Biological aging is the accumulation of damage to the cellular and molecular functional units of tissues over time, leading to breakdowns of physiological systems, age-related disease, and, ultimately, mortality.^1^ This process, combined with the global population undergoing a demographic transition in which a rising percentage is living into historically advanced ages, motivates the search for therapies that would intervene in the biological aging process and thus delay, decelerate, or reverse multiple manifestations of age-related ill health.^2^

The cornerstone test for longevity therapeutics in animal models is the lifespan study, emphasizing maximum lifespan.^3,4^ However, potential treatments should also increase the time spent in good health, motivating interest in developing quantitative metrics for health span. One approach to this metric is the rodent frailty index (FI). In humans, frailty is a clinical syndrome characterized by a decline in physiological resilience and increased vulnerability to adverse health outcomes, particularly in the elderly.^5^ As a population ages, there is an increase in frailty, especially over the age of 65, with a significant correlation between frailty, mortality, and hospitalization being evidenced.^6,7^ Hence, researchers have constructed practical and objective measures of frailty to manage and test new interventions against the syndrome. The frailty index is a quantitative, multidimensional approach that evaluates an individual’s level of frailty based on the accumulation of health deficits.^8^ Subsequent research has shown that FIs are robust predictors of mortality in humans, even in people early in middle age who are not ordinarily frail.^9,10^ FIs have been proposed as potential biomarkers of aging, and similar models are responsive to putative longevity interventions in humans.^10–13^

More recently, rodent FIs have been developed on principles like the human originals, using a panel of physical and/or clinical parameters to measure frailty. However, developing FIs in rodents is not straightforward. Researchers have adopted different approaches by varying the types and numbers of parameters to be included in the FI and cut-off points that classify an animal as frail. This variation results in each study having a different operational definition of frailty, which reduces the ability to compare findings across studies, thereby impeding progress toward longevity therapeutics. Even in humans, where sample sizes are larger and clinical data for FI construction is broader and more robust, different frailty instruments vary in which patients they identify as frail and are challenging to implement in clinical settings.^14^

This perspective highlights these differences and compares rodent and human frailty index scores. We summarise the published strategies for implementing frailty indexes, discuss their strengths and weaknesses, and implement a frailty index based on physical performance with our young and old mice. We conclude by making recommendations regarding FI use for studies of aging, age-related disease, and longevity therapeutics.

## 2. Materials and Methods

### Animals

Young (3-4 months), middle-aged (18 months), and old (28 months), mainly male C57BL/6 mice, were acquired from Charles River Laboratories (Wilmington, Massachusetts) (n = 3-7 per group). Young 3-month-old (n=5) mice were all male, 4-month-old mice included two males and two females, 18-month-old (n=5) were all male, and 28-month-old (n=4) were three males and one female. Mice were housed in a vivarium with *ad libitum* access to food (Teklad Global Soy protein-free, Envigo) and water (Aquavive Water, Innovive Inc). Cages were changed every 1-2 weeks depending on the number per cage, and documents were maintained under national and local regulations. All animal experiments in this study are approved by the Lifespan Research Institute IACUC committee following the “Guide for the Care and Use of Laboratory Animals” prepared by the Institute for Laboratory Animal Research, National Academy of Sciences.

### Grip Strength

C57BL/6 mice were used to establish baseline grip strength using the Grip Strength Meter (47200, Ugo Basile®). The mice were held by the mid-base of the tail, allowing their front paws to grip the bar. Once gripped, the mice were pulled back steadily, keeping them horizontal. The flat bar was used to measure the highest forelimb grip strength of the mice. To account for weight increases in older mice, results were normalised to weight. Each mouse was tested 4 times. The weakest measurement was removed and the average of the remaining scores was calculated.

### Open Field Testing

C57BL/6 mice were used to establish a baseline of mobility and behavior using open-field testing (Noldus Ethovision XT 17.5). Two arenas side by side (50x50cm) were set up with Basler GenICam (acA1300-60, Basler) above. Arena settings were utilized to establish each arena’s scale (50cm) and center and periphery. The periphery is defined as the outermost 10cm on either side of the center (30cm) of the arena. Each mouse began the experiment in the bottom left corner and was recorded for 10 minutes. Video analysis software (Ethovision XT 17.5) was used to track the mice using the center body point and calculate the total distance moved, percent of time spent in movement, average velocity, body elongation, and mobility. Between each trial, 70% ethanol was used to clean the arena to remove scent marks, and a 10-minute wait period was used to allow ethanol evaporation.

### Measures used to construct the Frailty Index (FI)

The factors from open-field testing used to comprise the frailty index are previously described by Whitehead et al., 2013 and Parks et al., 2012. They include the total distance moved (tracked by center-point over 10 minutes in cm), the maximum distance between two consecutive points in the tracks in cm (available in trial statistics of distance moved in the Ethovision analysis profile), total duration of movement (s), proportion of time spent moving (%), meander (change in direction per unit distance moved measured in degrees/cm from 0^◦^-180^◦^), average velocity (cm/s), rearing frequency (occurrence/time), and weight (g). For both Parks et al. (2012) and Whitehead et al. (2013), the open-field test duration was 10 minutes.

### Frailty Index Cutoff Points

The young, middle-aged, and old mice required reference values to determine frailty levels. For these, we included our 3-4-month-old mice and the reference values from Whitehead et al. (2013), which were males aged 5 months (n = 5), and Parks et al. (2012), which consisted of males and females aged 13.5 months (n=3 per sex). When using our reference values from 3-month-old male mice, all females from our cohort were removed. Our entire cohort was measured using Whitehead and Parks reference values. For each mouse, each factor of the FI was compared to the reference values. The frailty scores were applied if the measured score differed from the reference by at least 1SD (Standard Deviation). They were graded as follows: less than 1SD scored 0; values differed by ± 1 SD scored 0.25; values that differed by ± 2 SD scored 0.5; values differed by ± 3SD scored 0.75; values that differed by more than 3SD scored maximum frailty of 1. As in Parks ’ study, the sum of these scores for each factor was divided by the number of parameters (8) to produce a total frailty score for each mouse.

### Statistics

GraphPad Prism V10.1.0 software was used to perform statistical analysis. A two-way ANOVA analysis was used to see if two or more independent variables affected the dependent variable (multiple age groups in frailty scores or grip strength). To be considered statistically significant, the P-value must be ≤ 0.05 and was quantified in the following order. * = p ≤ 0.05, ** p = ≤ 0.01, *** = p ≤0.001, **** = p ≤0.0001. For each experiment and each condition, n ≥ 3.

### Study Selection Process

A rigorous literature review with a range of keywords was used to search peer-reviewed journals: frailty, frailty index, aging, mice, longevity, healthspan, rodents, and phenotype. Databases searched included Pubmed, EBSCOhost, Google Scholar, and Loughborough University Ex Libris. This method provided 18 peer-reviewed articles published between 2012 and 2023. Other papers were identified but not included as this perspective focuses on frailty indexes that are novel, modified, or further validate established versions.

**Table 1.**
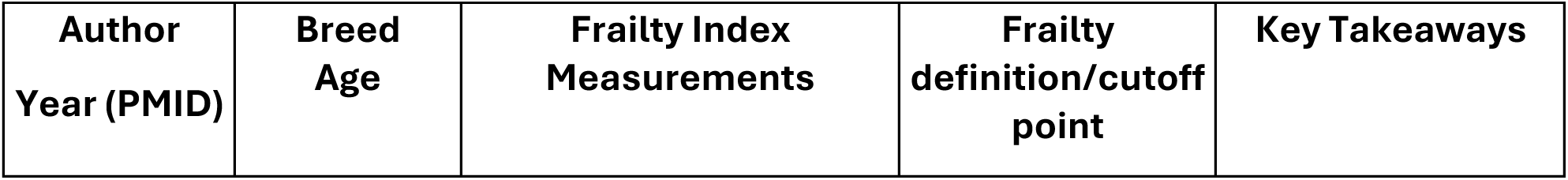
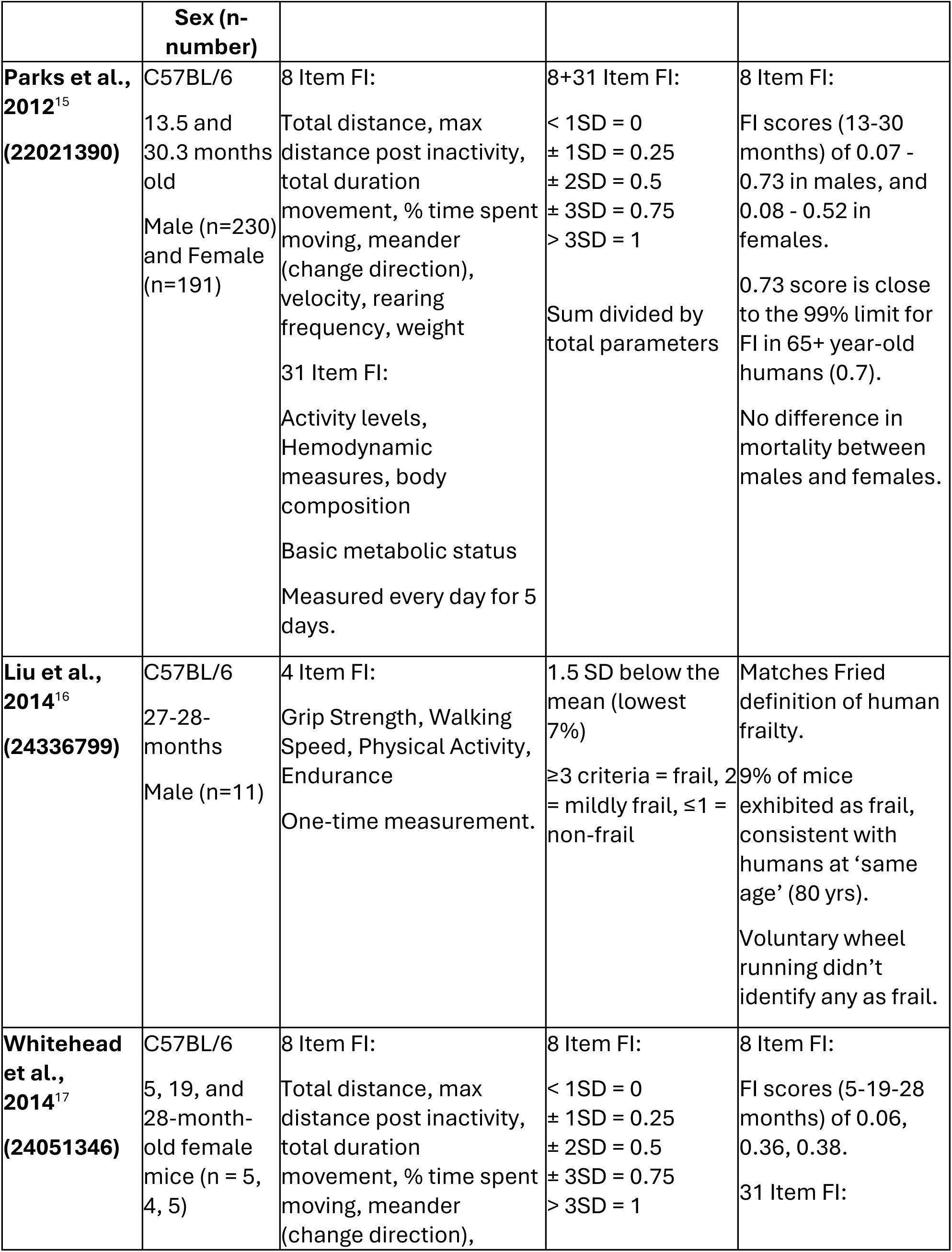
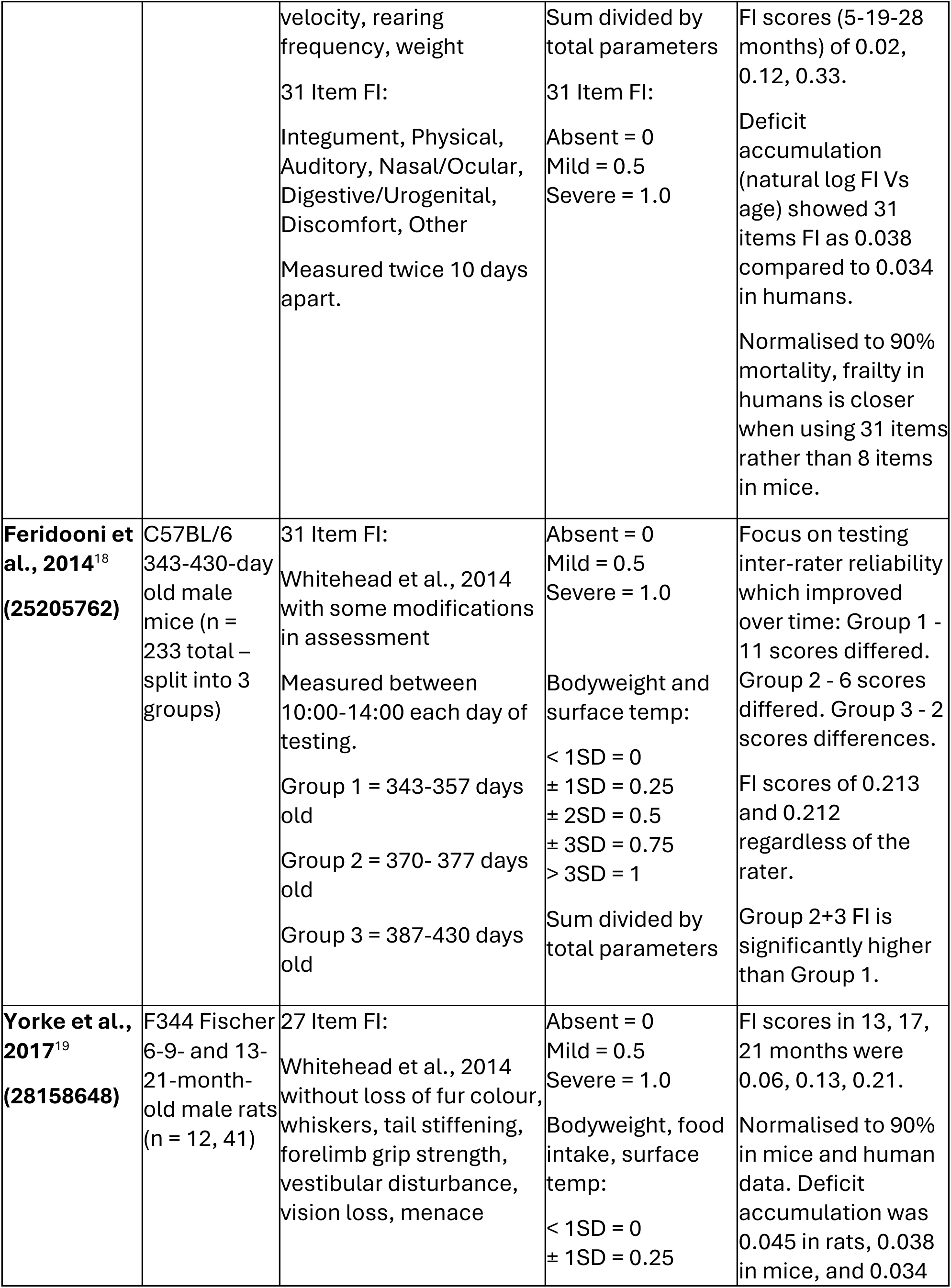
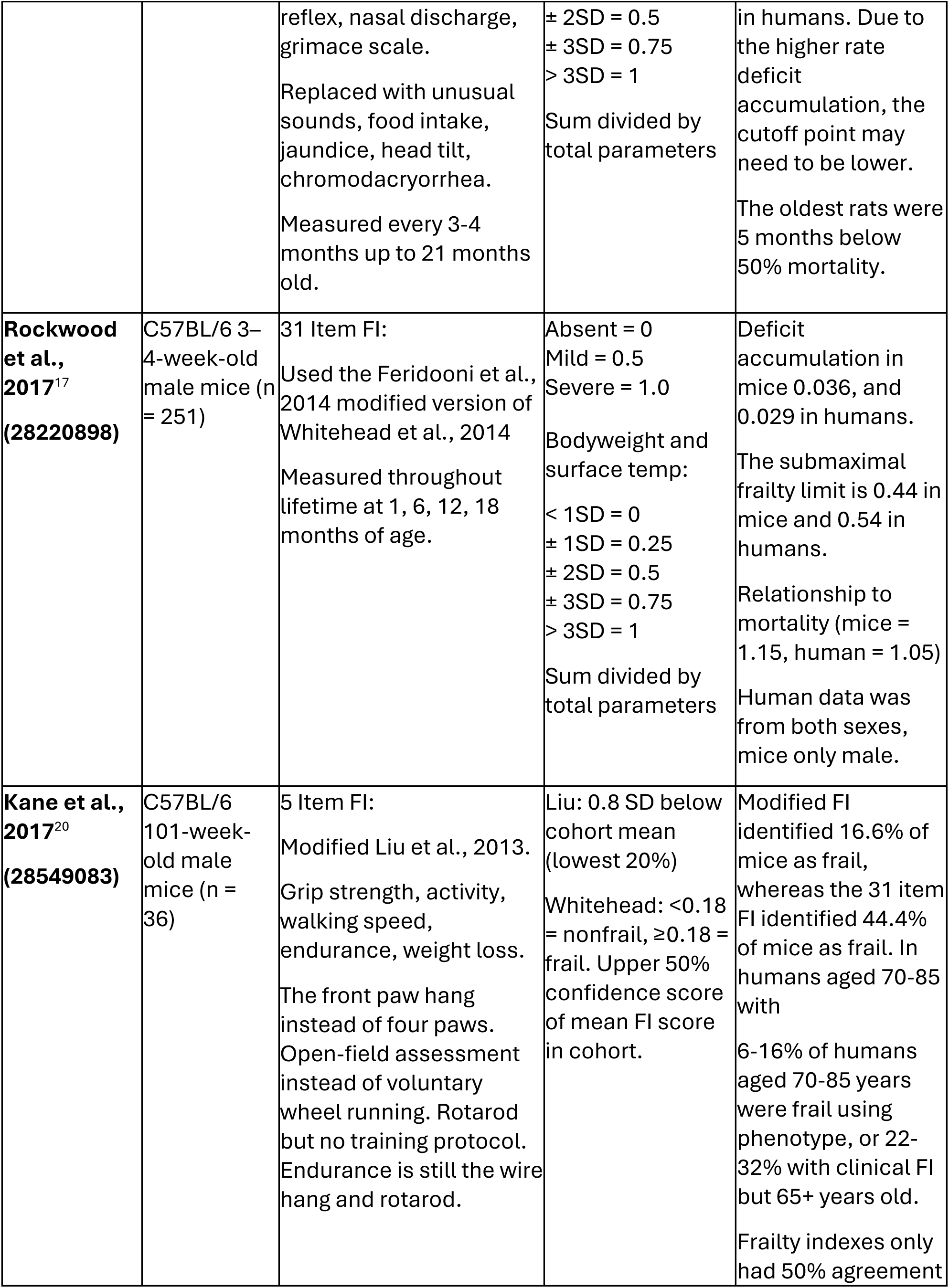
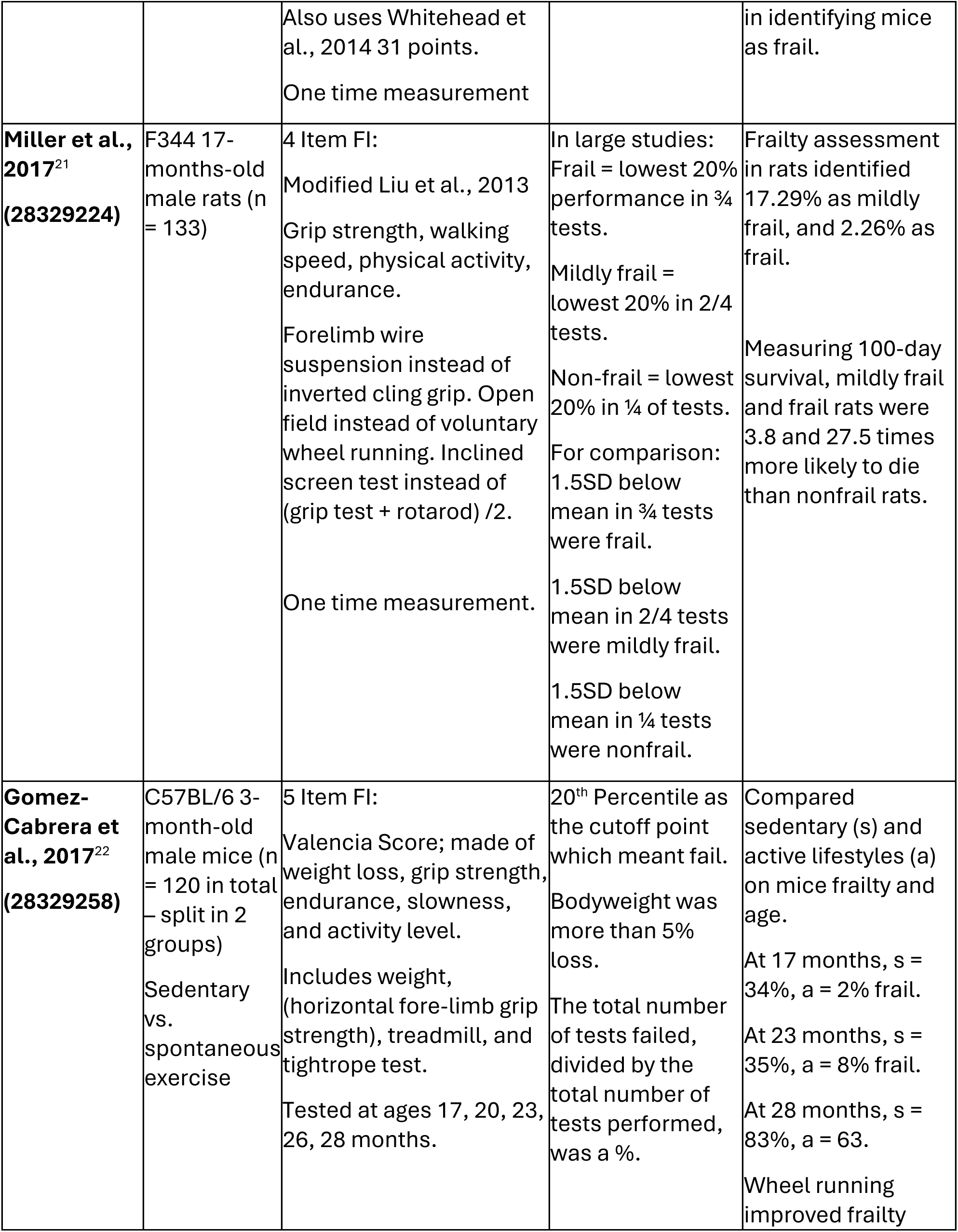
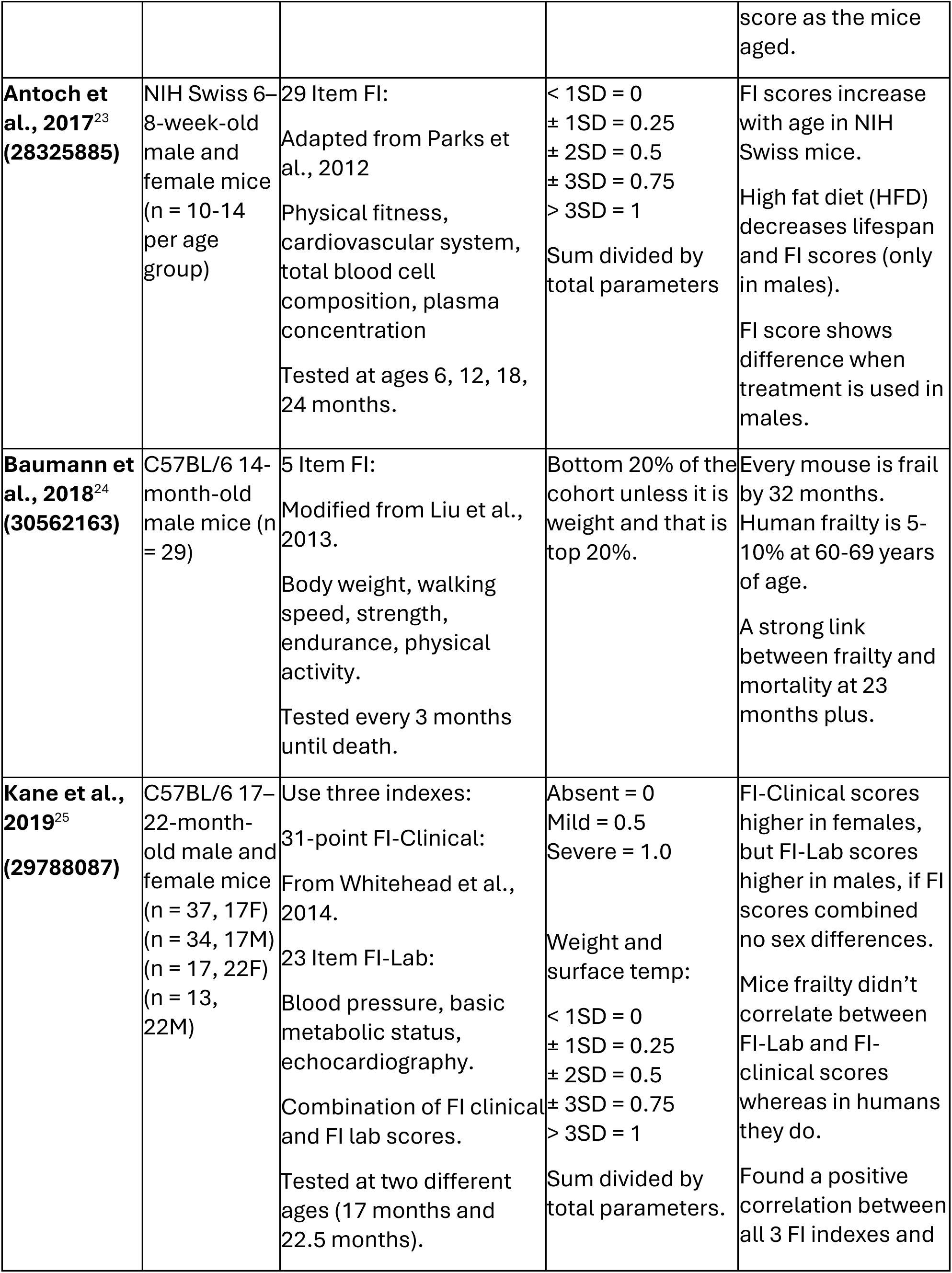
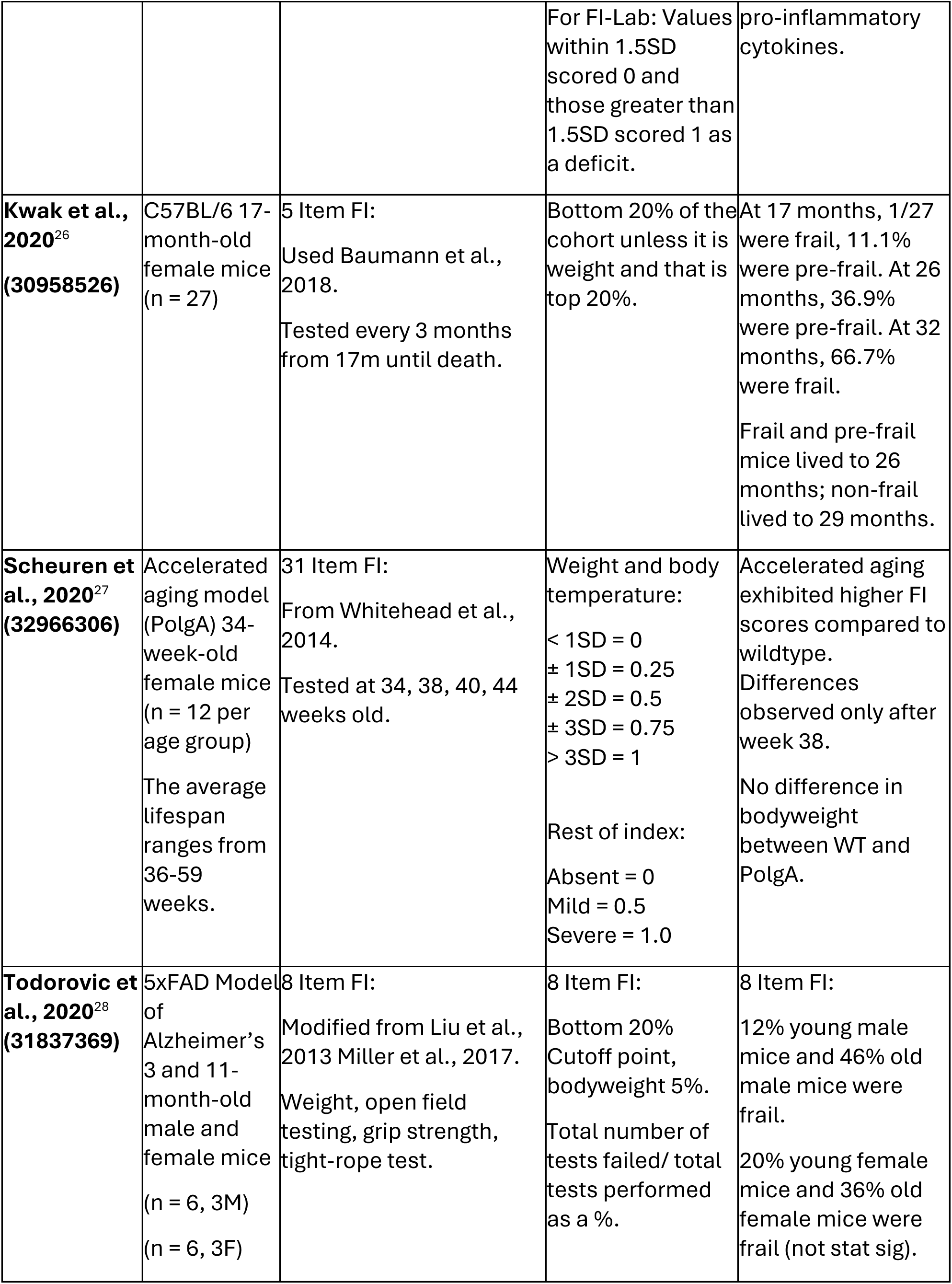
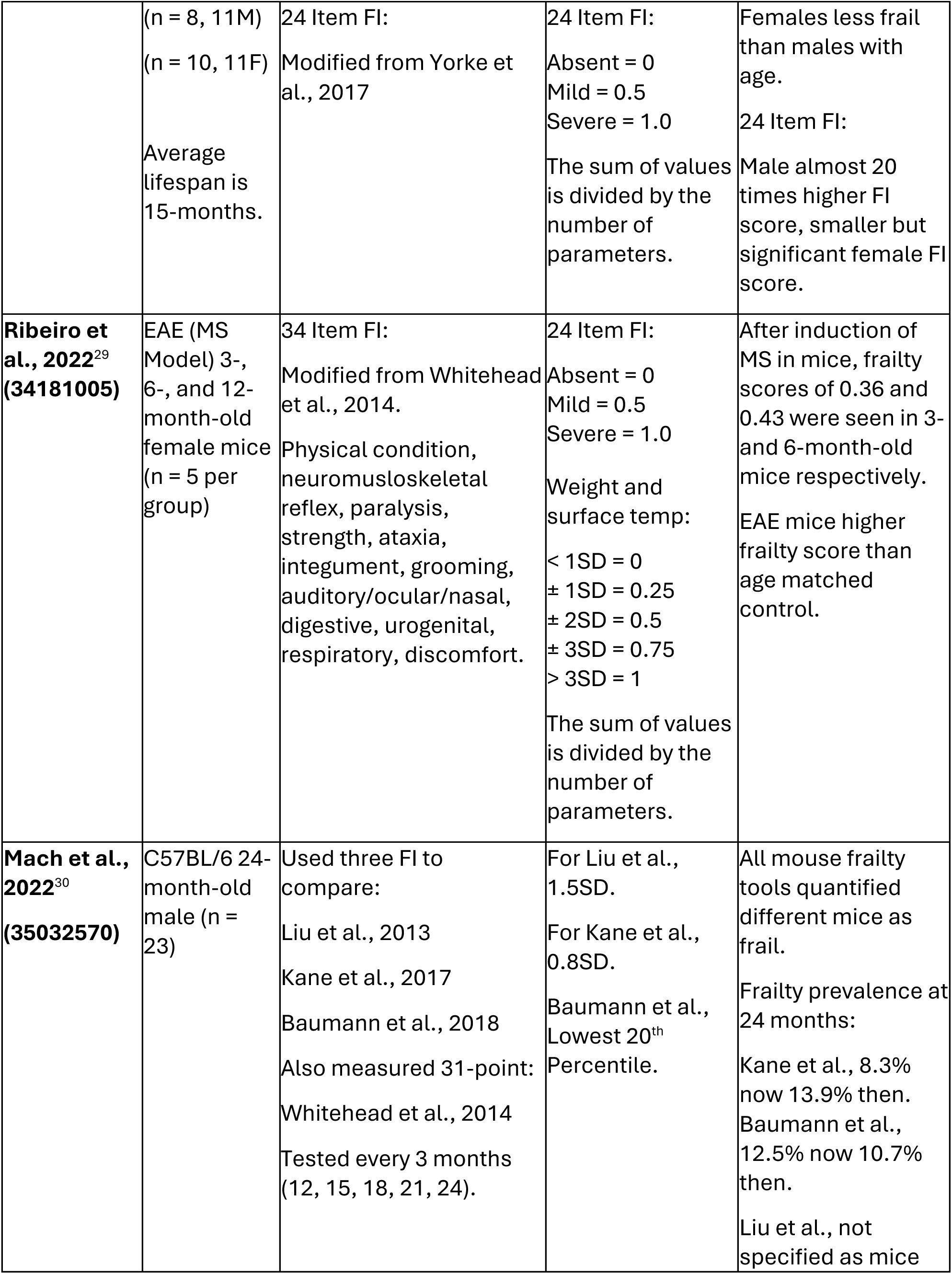
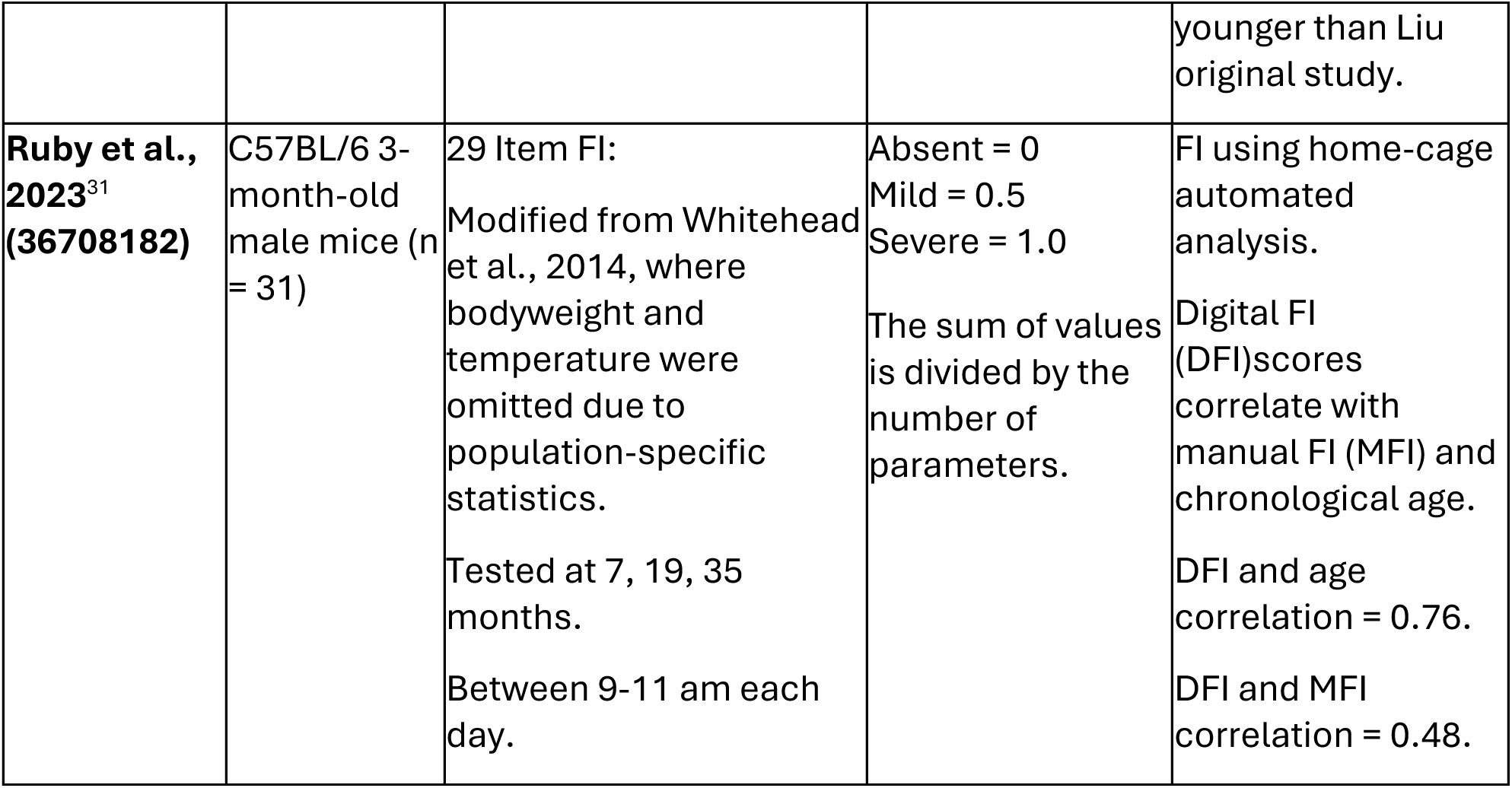
Summary of frailty index studies.

## 3. Results and Discussion

All FIs included in this review provide a numeric score indicating the level of frailty present, but their respective methods differ significantly. When choosing which established FI to implement, one must examine what cutoff point to utilize, what reference values to use if a scaled frailty score is desired or the cohort is small, the equipment available, and the factors to include. For instance, scoring systems may rely on quantifying physical performance (12/18 studies)^15,17,20–24,26,28,30–32^ or clinical observations (13/18 studies)^15,17–20,23,25,27–31,33^, with several articles using both.^15,17,20,23,28,30,31^ Those measuring physical fitness derive their basis from Fried et al., 2001, who created a human frailty index measuring four key factors: weakness, slowness, low activity, and poor endurance.^34^

The contents of the FIs vary significantly, depending on whether clinical observations or physical outputs are measured. For instance, those focusing on physical measurements have fewer items in their FI (4-5 items (7/18 studies)^16,20–22,24,26,30^ or 8 items (4/18 studies))^15,17,28,31^, whereas those using clinical observations measured 23-34 items (12/18 studies)^15,17–19,23,25,27–31,33^. Additionally, there is variation in the cutoff points used to determine whether a subject is frail and to what extent. This includes 0.8SD from a reference point or the lowest 20% of a cohort (7/18 studies)^20–22,24,26,28,30^, 1.5SD (4/18 studies)^16,21,25,30^, a staggered cutoff point of 1, 2, 3, 3+SD (9/18 studies)^15,17–19,23,25,27,29,33^, or visual determination of 0 = not frail, 0.5 = mildly frail, and 1 = frail (9/18 studies)^17–19,25,27–29,31,33^. The cohort mean is often utilized as the reference point. But if a staggered cutoff point is used, a reference value is required from a control subject group. If the cohort mean-SD is used as the cut-off point, the frailty scoring is binary, either not frail = 0 or frail = 1. 0. When multiple cut-off points are used (1, 2, 3, 3+SD), then frailty is scored in a gradient (0, 0.25, 0.5, 0.75, 1).

While human frailty is the basis for rodent FIs, how to compare data from rodent FIs to equivalent effects if translated to humans is not clear. One approach is to compare deficits among animals and humans of similar biological age. Various investigators have developed systematic methods of equating biological age between mice and humans, including development, epigenetic age clocks, gene expression patterns, disease onset ages, median and maximum lifespan proportions, and/or the trajectory of the survival curve.^35–38^ As a result, several studies ^15–17,19,20,24,25,33^ have linked their rodent analysis to quantitative human data, whereas others make no direct comparison ^18,21–23,26–31^.

Those making such comparisons often use deficit accumulation as the key metric (natural log of FI vs. age) after normalizing human and mouse data sets to 90% mortality values or comparing the equivalent ages of the two species. The problem with using corresponding ages is the inconsistency across the literature in what age cutoffs are considered equivalent. Liu et al. (2013) identified 9% of 27-28-month-old mice as frail, consistent with frailty levels in 80-year-old humans. Baumann et al. (2018) found all mice frail at 32 months of age but compared this to 60+ years in humans, where 5-10% of 60-69-year-olds or 26-65% of 85+- year-olds are frail. Another publication suggests that a 32-month-old mouse is equivalent to a 109-year-old human.^39^ Kane et al. (2017) observed 16-44% of 23-month-old mice as frail depending on the FI index used, while humans aged 65+ years showed a 22-32% frailty range using comparable indexes. Two FIs quantified different animals of the same age group as frail.^20^ Furthermore, another study compared three FIs in a group of 24-month-old males and found inconsistency between which mice were frail.^30^

Like the heterogeneity of mouse data, the lack of a universally agreed standard of human frailty scoring ensures that the reference ages for the percentage of the population identified as frail at a given age will also be inconsistent. The difficulty in making precise comparisons is evident, though the underlying fact of deficit accumulation increasing with age remains. Developing a simplified approach for FI calculations in murine models and humans is essential for standardizing assessments, improving reproducibility, and validating medical interventions for their translational potential for human aging and frailty management.

We strived to provide recommendations for implementing a FI in murine models with commonly available equipment and cohort size in typical studies. To this end, we scored our mice on the 8-item FI developed by Parks et al. (2012) and implemented by Whitehead et al. (2014), as they suited the available equipment and allowed frailty to be measured as a gradient with relatively few animals. To score the mice in our study, we utilized the reference values from these studies. As the reference values of Parks had both sexes and our study did too, they were sex matched. Notably, the Whitehead cohort was female, and most of our subjects were male. However, the 3-4-month-old mice in this study are younger than those used to establish the reference values for both Parks and Whitehead.

Using Parks et al. (2012) reference values, the 8-item FI yielded frailty scores of 0.37/1 for our 3-4-month-old mice, 0.52/1 for our 18-month-old mice, and 0.66/1 for our 28-month-old mice (Figure 1.A.). Using Whitehead et al. (2014), reference values (averages of trial 1+2) identified the 3-4 months with 0.26/1, 18 months as 0.46/1, and 28 months as 0.48/1 frailty scores (Figure 1.A-i). In both assessments, the frailty scores among the 3–4-month-old animals were notably high. The parameters responsible for scoring young mice as frail were the maximum distance post inactivity, meander, and movement duration, which affected this group’s overall frailty score.

**Figure 1.**
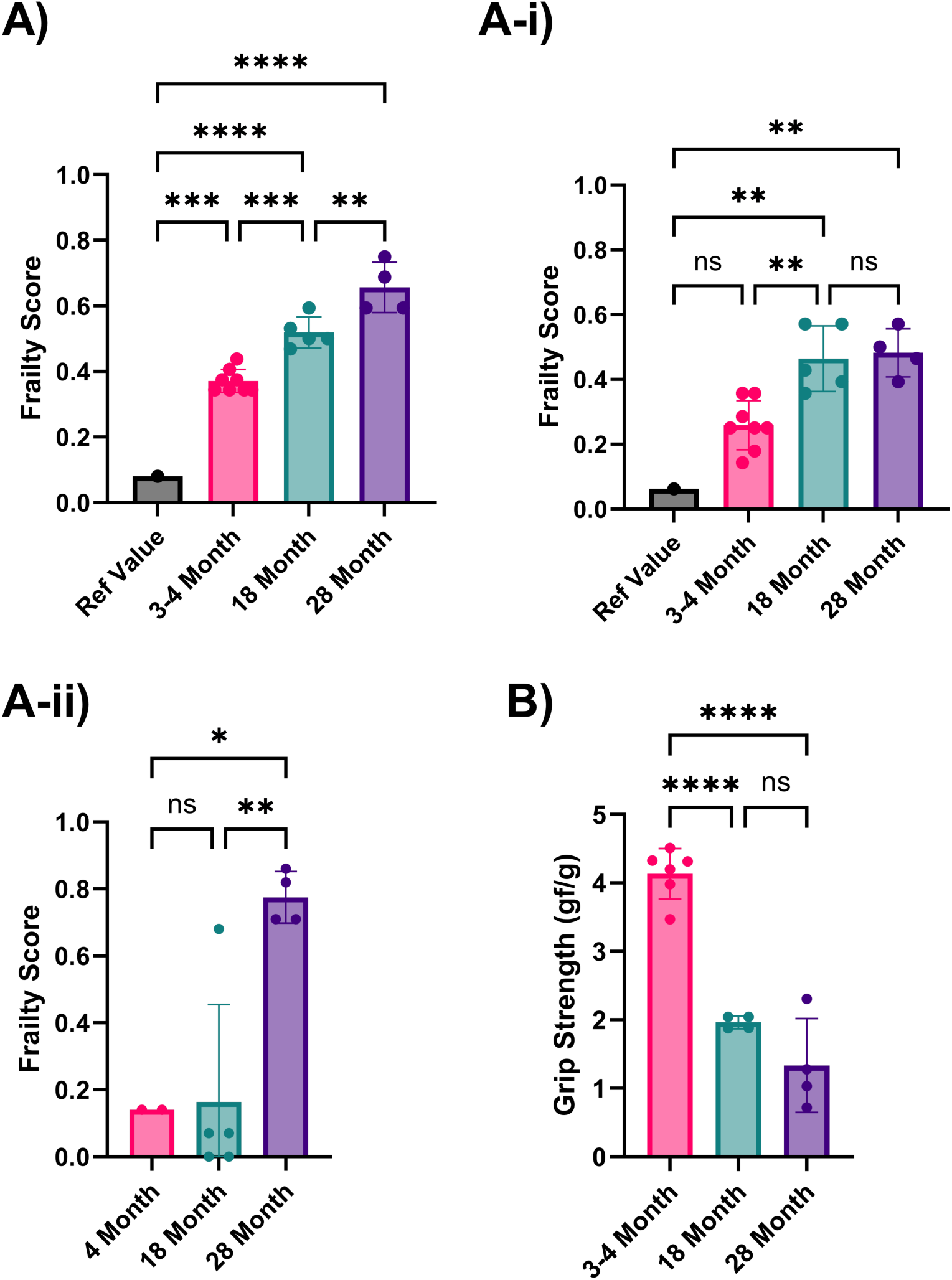
Analysis of the frailty phenotype in 3-4, 18, and 28-month-old C57BL/6 mice. A) Frailty index scores implemented using Parks et al., 2012 reference values. A-i) Frailty index scores implemented using Whitehead et al., 2014 reference values. A-ii) Frailty index scores using our own 3–4-month-old mice as reference values. B) Grip strength (average forelimb parallel bar score) is normalized to weight for 3-4, 18, and 28-month-old mice. P-values ≤ 0.05 (*), ≤ 0.01 (**), ≤ 0.001 (***), ≤ 0.0001 (****) were calculated using one-way ANOVA for three or more independent groups or unpaired t-tests for just two independent groups. Columns represent the mean with error bars ± SD.

High scores in movement duration in our young mice may be due to differences in acclimatization protocols. Parks et al., 2012 allowed 5 days of testing in the arena, using the last 2 days for assessment. We acclimatized the mice to the testing room but not the arena for an hour before data collection, as open field testing depends on the inherent explorative nature of mice. The meander can also be measured differently, either as a relative or absolute meander, using Ethovision video software analysis (version 17.0.1). It can also be calculated from either the body point or head direction. Maximal distance post inactivity could also use further clarification on how it is precisely measured; for example, how inactivity was defined, and what body point was measured.

Based on these potential sources of discrepancy, we recommend each lab use its own reference mice to limit variability. When we did this using our 3-month-old mice to establish the reference values, the overall frailty scores in mice aged 18-28 months were much lower than when Parks et al.’s or Whitehead et al.’s reference values were used (0.03/1 and 0.22/1, respectively (Figure 1. Aii)). Most studies (12/18) have used C57BL/6 mice to assess new FIs or modify existing FIs. Therefore, if using a different mouse strain or a transgenic or disease model, identifying the baseline of a lab’s rodent population is of yet further importance. We recommend using each mouse as its reference point for medical intervention or longitudinal studies, strengthening the analysis without increasing the workload.

One challenge when creating reference values from a group of young mice is inherent variation. If the reference group variation is significant, some of the frailty parameters in test mice cannot reach the score of 1. One strategy to circumvent this is to reduce diversity in the reference value group. For example, Antoch et al. (2017) excluded animals if their scores exceeded by more than one SD from the mean. However, including or excluding outliers in small sample sizes risks creating non-representative values. Using each mouse as its reference point also circumnavigates the potential issue. Similarly, while FIs can separate young and old mice, it is less certain they can determine the difference between a treated and untreated group of the same age. To ascertain if a treatment can improve frailty, we suggest scoring the same mice before treatment and utilizing that as reference values, rather than comparing old, treated mice to younger untreated mice as commonly done in rodent aging studies.

Akin to 4-5 item FIs, we included grip strength to characterize the physical health of the mice further. Due to equipment limitations, we could not fully implement the 4-5 item FI which required an inverted cling-grip test, rotarod, and voluntary wheel running cages. In grip strength scores normalized by weight, the 3-4-month-old mice had the highest average score, with 18-month and 28-month-old mice having consequently lower scores. However, the difference between 18 and 28 months was not statistically significant due to a single animal’s exceptionally high score (Figure 1B). Using individual mice as their comparators in longitudinal assessments could help identify any increases or decreases in performance. One limitation of this study is the low number of animals per group, which limits the robustness of results for animals of a given age. However, this perspective may help in frailty-scoring selections for small-scale studies with limited subjects.

Open-field testing (OFT) or automated video testing simplifies frailty measurements when the methods are clearly defined. Furthermore, it eliminates errors introduced by limited inter-rater reliability, a well-recognized problem in clinical observation FIs, as user input is minimal.^18,19^ To determine the best factors from OFT to include, an independent study is required to determine which parameters best predict mortality. For instance, investigators could make periodic measurements of the various OFT parameters (e.g., every 3 months) until death, then assess each factor to determine its weight for predicting mortality compared to an established FI. Principal component analysis should be applied to reduce the number of variables and choose those that are both highly predictive of mortality and (amongst those that fall within the same principal component) most convenient for implementation.

The reason for this extensive independent study is the components of current indexes. For instance, 8 item FIs include the total distance (cm), velocity (cm/s), and movement duration (s and %). Therefore, 4/8 factors measure closely related physiological traits, giving movement great weight in this index. Although walking speed is a well-established mortality indicator and predictor of surgical outcomes in humans, assigning movement-related variables half the overall score in a rodent FI will likely overweight the index.^6,7^ One study prioritized a diversity of health-related physiological systems over one key physiological trait, alongside quantitative parameters without visual scoring while being minimally invasive.^23^

For future additions to FIs, one study measured gait speed in the cage and on the wheel in C57BL/6 mice and found it correlated with age and the manual frailty index.^31^

## 4. Conclusion

Studies discussed in this perspective offer a variety of approaches to measuring frailty. We recommend that investigators carefully consider what aspects of frailty to include in their analyses instead of fully adopting the published scoring systems. It is preferable to avoid including and giving equal weight to multiple parameters that measure closely related physiological traits, such as multiple parameters related to movement. In addition, it is best to use automated measurements where possible to avoid experimenter bias and inter-rater variations. As there is substantial variation even between mice of the same strain, it is optimal to measure baseline frailty in each animal and to track changes over time in response to treatment and during physiological aging, especially in a small sample size. Doing so accounts for variability in baseline values. It increases sensitivity in detecting subsequent changes while reducing the chances of false positives, resulting in a more reliable measurement of physical health in aging rodents and the effects of longevity therapeutic candidates.

## Author Contributions

O.G.F. analyzed the literature, generated the data, FI scores, table, and graph, and wrote the manuscript. A.B. generated data and edited the manuscript. M.R. edited the manuscript.

M.A. helped generate data. A.R. conceptualized the frailty measurement project, supervised its execution, and edited the manuscript. A.S. conceptualized the manuscript, supervised data generation, and edited the manuscript.

### Funding support

We want to acknowledge funding agencies who supported this work at the Longevity Research Institute and Loughborough University (EPSRC).

### Conflict of interest

The authors declare no financial interests related to this work.

## Notes

### Competing Interest Statement

The authors have declared no competing interest.

